# Distinct effects of priming the brain using tDCS and observational practice: new evidence from brain effective connectivity

**DOI:** 10.1101/2023.12.12.570884

**Authors:** J. McLeod, A. Chavan, H. Lee, S. Sattari, S. Kurry, M. Wake, Z. Janmohamed, N.J. Hodges, N. Virji-Babul

## Abstract

Complex motor skills can be acquired while observing a model without physical practice. Transcranial direct-current stimulation (tDCS) applied to the primary motor cortex (M1) also facilitates motor learning. However, the effectiveness of observational practice for bimanual coordination skills is debated and there is little research on the effects of tDCS on acquiring bimanual skills and the underlying effective/causal brain connectivity. We compared the effect of primary motor cortex tDCS (M1-tDCS) to action-observation (AO) when acquiring a bimanual, two-ball juggling skill and characterized the brain causal connectivity patterns underlying each condition. Twenty healthy young adults with no juggling experience were randomly assigned to either video observation of a skilled juggler or anodal M1-tDCS. Thirty trials of juggling were performed and scored after the intervention. Resting-state EEG data were collected before and after the intervention. Information flow rate was applied to EEG source data to measure causal connectivity. Juggling scores were significantly higher in the AO group (*p* =.03). We found the strongest information exchange from (L) parietal to (R) parietal regions, strong bidirectional information exchange between (R) parietal and (R) occipital regions and an extensive network of activity that was (L) lateralized in the AO condition. In contrast, the M1-tDCS condition was characterized by bilateral long-range connections with the strongest information exchange from the (R) occipital region to the (R) temporal and (L) occipital regions. This study provides new results about the distinct network dynamics of priming the brain for skill acquisition using direct stimulation or indirect stimulation via action observation.

## 1. Introduction

Motor learning can be facilitated without physical practice by repeatedly observing the performance of a motor skill, referred to as observational learning (Hodges, 2017; Hodges et al., 2007; Ramsey et al., 2021). Observational learning of motor skills is thought to involve a complex interaction within a distributed brain network involving motor as well as non-motor regions including the inferior parietal cortex (IPC), superior temporal sulcus (STS), inferior and superior frontal cortices and occipito-temporal regions with mirror-like properties that respond both to the execution and observation of actions (e.g., Calvo-Merino et al., 2006; Grèzes et al., 2003; Virji-Babul et al., 2009; see Ramsey et al., 2021 for a recent review). Much of the research in observational learning is on the adaptation or adjustment of actions that can already be performed, such as adaptation of throwing actions to a target, pressing keys to memorize a sequence, or tracking tasks. In some of our own research involving repeated observation of a joystick tracking task, observational practice induced neurophysiological changes as indexed by mu suppression at central sites, providing further evidence for motor-based processes of the AON being active during observational practice. However, we did not find that these motor-related processes were related to behavioural measures of learning (Alhajri et al., 2018). In addition to the need to establish direct links between measures of brain activity and behavioural outcomes, little is known about the brain processes underpinning observational learning of new actions, when the observer does not have an existing motor skill set to perform what they are watching.

Transcranial direct current stimulation (tDCS) modulates cortical excitability and reorganization through the application of a weak current through the skull (Nitsche & Paulus, 2000, 2001). Over the past decade, there have been numerous reports evaluating the effects of tDCS on human motor performance and learning. Increasing the excitability of the primary motor cortex (M1) with tDCS during motor training has been shown to improve motor performance for a range of skills such as the sequential finger tapping task (Karok & Witney, 2013; Rumpf et al., 2017; Stagg et al., 2011), serial response reaction time tests (Ehsani et al., 2016), and visuomotor tracking (Goodwill et al., 2013). These effects have primarily been reported for unimanual tasks (see Pixa & Pollok, 2018 for a review) with few-to-none evaluating the uncoupled effects of tDCS. Despite the limited number of studies investigating bimanual motor skills, Pixa and Pollock (2018) suggest that tDCS has the potential to enhance bimanual motor performance.

Effective connectivity (EC) provides a measure of the influence (direct or indirect) that one brain region exerts over another (Friston, 2011) and identifies causal, directionally-dependent interactions between different brain regions. Little is known about the effects of tDCS at this network level, particularly when tDCS is applied over the motor network. Calzolari et al. (2023) evaluated resting state effective connectivity across the motor network after applying tDCS over M1 or the cerebellum in the same cohort on separate days. They reported changes beyond the targeted regions that were stimulated, including between the cortex, thalamus and cerebellum.

However, they focused their analysis on the resting state activity of the brain without using a motor learning task. A key question that arises from their work, and observational learning research in general, is how is causal connectivity between brain regions modulated during action observation as compared with the application of M1-tDCS, when acquiring a novel bimanual task? In both cases, there is expected to be activity in motor-related regions through direct or indirect stimulation of the motor cortex and surrounding regions.

We investigated the patterns and statistics of effective connectivity using a data-driven effective brain connectivity measure that is based on the concept of information flow rate applied to EEG signals (see Hristopulos et al., 2019). The information flow rate was developed by Liang using the concept of information entropy and the theory of dynamical systems (Liang, 2013). Information entropy is a measure of the information contained in a given signal (e.g., time series) and quantifies the changes in the information content of a time series (and hence, the temporal evolution of the brain region from which the signal is acquired) as a result of the interactions with other brain regions and the stochastic forces. The information flow rate measures the directional transfer of information between time series at different locations and therefore between different brain regions. A high information rate from region A to region B suggests that a large amount of information is transferred from A to B per second. Given a collection of source locations in the brain, the information flow rate can identify which sources transmit and which receive information, thus leading to a network of brain connections. Since the derived connectivity is directional, the information flow rate provides a robust method for detecting causal links in the brain.

The aim of this study was to use information flow rate to measure effective connectivity following AO and M1-tDCS in two groups of healthy individuals. Such measures would allow us to compare the brain activity associated with direct and “indirect” stimulation (through passive observation) of motor-related brain areas, detailing the direction and spatial patterns of information flow. We were also interested in examining the outcomes of skill acquisition of a novel bimanual task resulting from the pairing of AO and M1-tDCS separately. We chose juggling as the novel bimanual coordination task, which requires simultaneous control and coordination of multiple movements (Berchicci et al., 2017).

Given the limited previous literature related to effective connectivity on observational practice and tDCS on novel bimanual tasks, our preliminary hypothesis extended from past research on unimanual tasks. We predicted that: a) AO would be associated with activity in bilateral motor regions embedded within connections of the frontal-temporal-parietal action observation network and b), M1-tDCS would be associated with a dominant nexus of information flow arising primarily from the (L) primary motor region with bidirectional information exchange over widespread cortical connections.

## 2. Materials & methods

### 2.1 Participants

Twenty participants with no juggling experience from the University of British Columbia were recruited for a study on the effects of tDCS and AO. Participants were randomly allocated to one of two intervention groups; Action Observation (AO) or M1-tDCS (mean age: 23 ± 3 y*r*; 13 F; 7 M). Inclusion criteria included right-handed (based on the Edinburgh handedness inventory, Oldfield, 1971), normal or corrected-to-normal vision, no history of neurological or psychiatric disorders nor any contra-indications for tDCS (i.e., history of seizures or an intracranial implant). This study was approved by the University of British Columbia Research Ethics’ Board-B (H17-03361). Consent was obtained from all participants before enrolment.

### 2.2 Study Design

The chronology of the study is outlined in Fig. 1. All participants completed two, pre-intervention juggling trials, which were video-taped and saved for later analysis. A three-minute, eyes-open baseline EEG session was then recorded after which participants were randomly assigned to one of two fifteen-minute intervention groups: (1) video observation of a skilled juggler/ “AO group” or (2) M1-tDCS with no video observation/ “M1-tDCS group”. There was then a second three-minute, eyes-open resting state post-intervention EEG period, immediately followed by thirty practice attempts at the two-ball juggling task, which were video-taped for analysis.

**Fig. 1.**
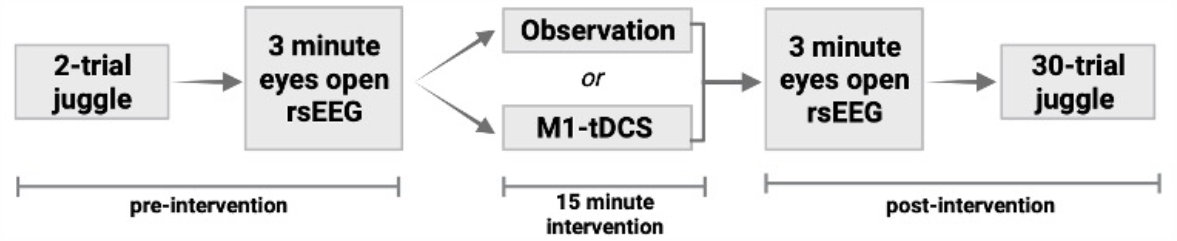
Experimental procedure. All participants performed a 2-trial juggle followed by 3-minute eyes open resting state EEG (rsEEG). Participants were randomized to one of two interventions: observation or tDCS stimulation. Individuals in the observation group watched a video of a skilled juggler for 15 minutes. Individuals in the tDCS group received 15 minutes of weak direct current (2mA) of anodal stimulation over the primary motor cortex (M1-tDCS).

### 2.3 Juggling Task

Effects of the practice intervention were investigated using a two-handed “exchange” juggling task with two balls (Zentgraf & Munzert, 2009). One juggling trial consisted of two throws and two catches and participants were instructed that the balls had to cross in the air and that they had to toss the balls to the opposite hand. The two juggling balls were identical in size and weight. Participants were seated while attempting to juggle. A performance analysis was subsequently conducted on the video recordings according to scoring criteria used by Hodges and Coppola (2015). Performance was scored using three criteria: hand release asymmetry, ball height symmetry, and whether one or two balls were caught. Each of the three criteria received a score ranging from 0 (symmetrical release, hand-over, low height, no balls caught) to 2 (asymmetrical release, approximately symmetrical peaks and both balls caught). The scores for each criterion were summed to obtain a total score for each juggle, with six denoting a perfect score. Two raters independently scored a subset of the data and agreed on the performance criteria using a standard process of consensus scoring.

### 2.4 Intervention

#### 2.4.1 Video Observation

Participants assigned to the video observation-only group watched a 15-minute video of a skilled juggler performing the two-ball juggling task from a 3^rd^-person perspective (Fig. 1C). Participants were not permitted to imitate the juggler while watching the video.

#### 2.4.2 M1-tDCS

Participants assigned to the tDCS-only group, received 15 minutes (2mA) of anodal stimulation (Starstim Neuroelectrics, Barcelona, SP) over the left primary motor cortex (M1) according to the international 10–20 electroencephalogram (EEG) system. Participants were asked to keep their eyes open during the stimulation.

### 2.5 EEG recording and preprocessing

Resting-state brain activity was recorded with electroencephalography (EEG), using a 64-channel HydroGel Geodesic Sensor Net at a 500 Hz sampling rate. Scalp electrode impedance values were confirmed to be below 50k before recording. All recorded signals were referenced to Cz. Raw EEG data were preprocessed using EEGLAB in MatLab. Each subjects’ EEG time series was re-referenced to the average, down-sampled 250 Hz, and notch filtered at 60Hz. We then applied a low-pass filter at 50 Hz and a high-pass filter at 0.5 Hz. Independent Component Analysis (ICA) was used to identify and remove any non-brain artifacts. Clean EEG sensor level data was converted to source space using Brainstorm in MatLab (Tadel et al., 2011). An inverse modelling method of minimum norm estimate (MNE) with sLORETA was used. The ICBM152 template was used as the head model. The source space solution was projected to ten regions of interest (ROIs) based on the Deskan-Killiany cortical atlas.

### 2.7 Effective connectivity

A full description of the information flow rate for application in the analysis of EEG source-reconstructed signals is provided in Hristopulos et al. (2019). Below we provide a summary of the methodology:

The information flow rate measures the rate of information transfer from the time series i to the time series j, and can be expressed using the above definitions as follows (Liang, 2014):

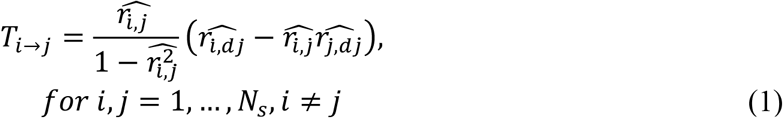

We refer to pi as the transmitter series and to pj as the receiver series with respect to Ti→j. A positive (negative) rate of information flow from i → j (Ti→j) indicates that the interaction between the two series leads to an increase (decrease) in the entropy of the series pj. It also signifies that the receiver series becomes more (less) unpredictable due to its interaction with the transmitter series. The predictability of each time series is negatively correlated with the entropy.

The information flow rate Ti→j is a measure of the information flow from series pi to series pj but it gives no indication of whether the impact of pi on the predictability of pj is significant. Quantifying the latter requires knowing the relative impact of the entropy transferred to the receiver from the transmitter series, compared to the total entropy rate of change due to all the influences acting on the receiver. The latter (hereafter referred to as the normalization factor for the information flow rate from pi to pj and denoted as Zi→j) can be computed (Hristopulos et al., 2019; Liang, 2014). The relative impact of the transmitter series on the receiver series is then given by the normalized information flow rate from the transmitter pi to the receiver pj: τi→j = Ti→j/Zi→j, which measures the percentage of the total entropy rate of change for pj that is due to its interaction with pi. Thus, in the following, we use τi→j to quantify the resting-state effective connectivity and use this measure to investigate the patterns of directional information flow between the different regions of the brain.

## 3 Results

### 3.1 Juggling Scores

Fig. 2 shows the averaged juggling scores for all participants pre-intervention (i.e., 2-trial juggle) and for each of the groups post-intervention (i.e., 30-trial juggle). Post-intervention juggling accuracy was significantly higher in the observation group compared to the tDCS group, t(18) = 2.30, *p* = .03, two-tailed (*d* =1.02).

**Fig. 2.**
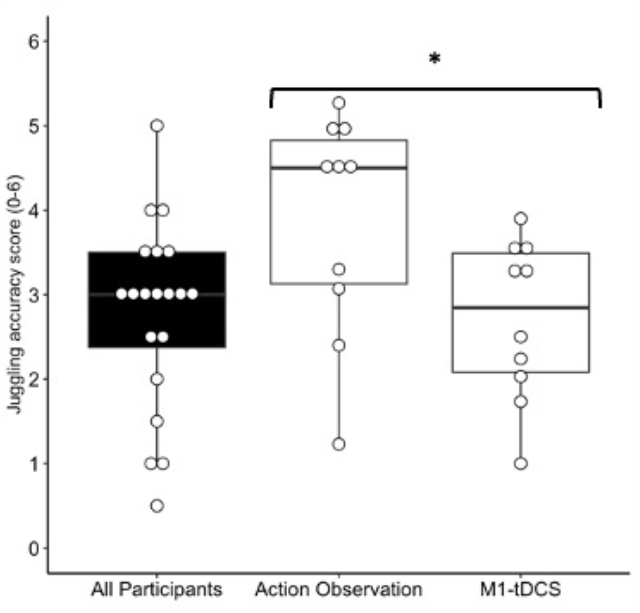
Mean total juggling accuracy score. Bars represent group mean and individual points represent participant scores. Error bars denote standard deviation. A two-sample t-test showed a statistically significant difference in post-intervention juggling accuracy between AO (M=3.88, SD-1.32) and M1-tDCS (M=2.71, SD=0.95).

### 3.3 Qualitative comparison of effective connectivity patterns

Based on the analysis of the EEG data, two individuals were excluded from the study due to excessive EEG noise making their data unusable, such that the final sample for the effective connecting analysis was n=10 in the AO group and n=8 in the M1-tDCS group.

Fig. 3A shows maps of the mean of absolute normalized information flow rate |*τi*→*j*| with transmitting regions on the y-axis and receiving regions on the x-axis and in 3B, maps of the top 10 mean |*τi*→*j*| values are shown. The source region labels are defined in Table 1. We limited the between-group comparison to the top 10 connections for the sake of simplicity. The top 10 |*τi*→*j*| values in the M1-tDCS group ranged from 0.034 to 0.078, and the AO group ranged from 0.031 to 0.051. The mean absolute normalized information flow rates |*τi*→*j*| are highest in the post-intervention M1-tDCS group compared to the AO group and pre-intervention baseline.

**Fig. 3.**
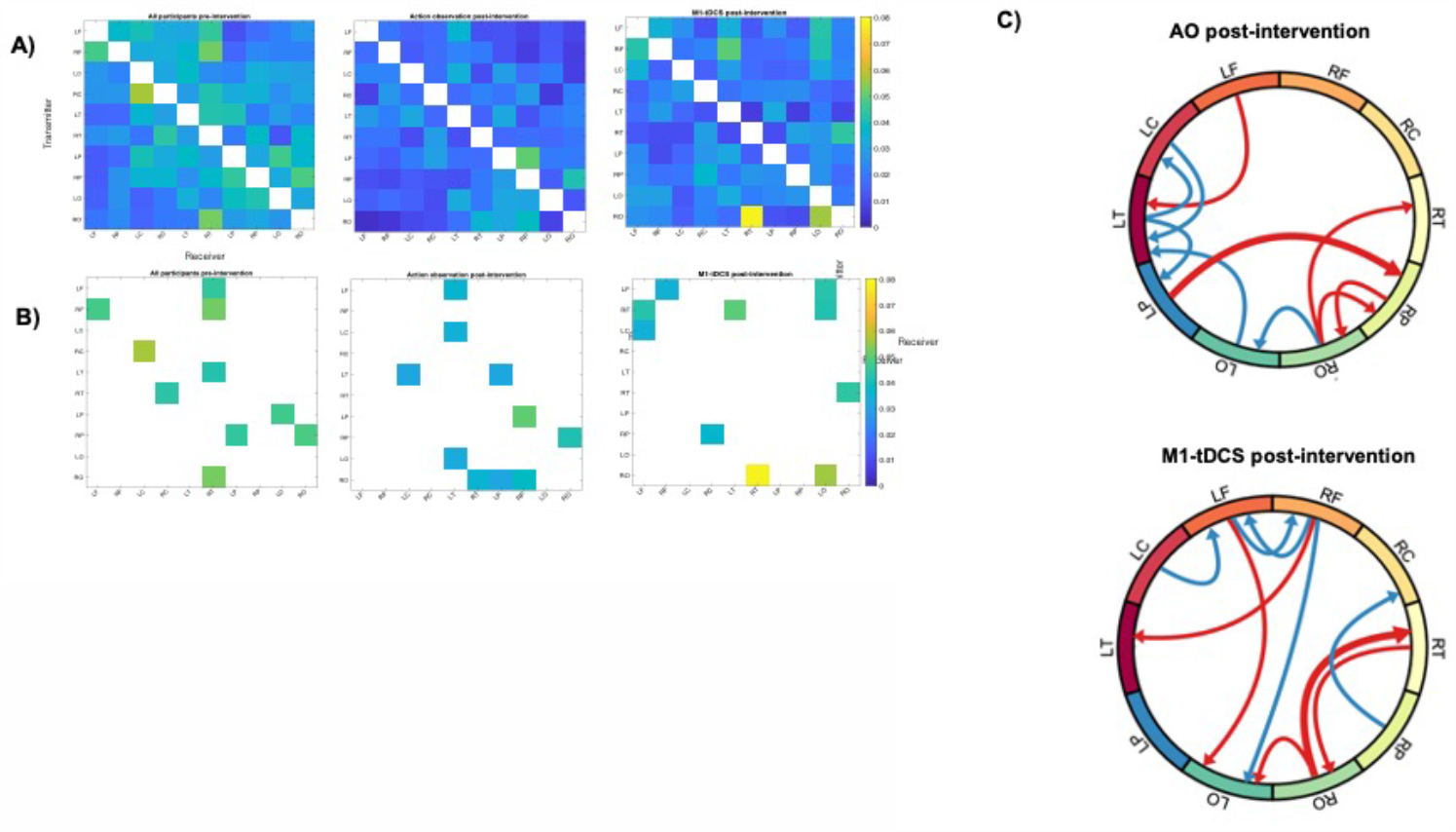
Causal connectivity analysis of EEG source data. Labels are listed in Table 1. **(A)** Maps of mean of absolute normalized information flow rate |*τi*→*j* |.Transmitting regions listed on the y-axis and receiving regions listed on the x-axis. **(B)** Maps of the top 10 mean |*τi*→*j* | values. **(C)** Qualitative summary illustration of the spatial distribution the connections with directional arrows for each group. The Illustration displays the top 10 connections ranked based on the average value of |*τi*→*j* |. The 5 most important connections are shown in red. The next five top connections (i.e. 6-10) are shown in blue.

Differences in the spatial distribution of normalized information flow rates |*τi*→*j*| pre-intervention and in the two groups post-intervention were seen (see Fig. 3B). In the AO condition, the strongest information exchange was from (L) parietal to the (R) parietal region, as well as strong bidirectional information exchange between (R) parietal and (R) occipital regions. In addition, there was an extensive network of activity involving the (L) frontal regions and bidirectional information exchange between central, temporal and occipital regions that was (L) lateralized. In contrast, in the M1-tDCS condition, information exchange was characterized by bilateral long-range connections, with the strongest information exchange from the (R) occipital region to the (R) temporal and (L) occipital regions. In addition, there were extensive long-range connections from (bi) frontal regions to (L) occipital regions and inter-hemispheric, bidirectional information transfer between the (L) and (R) frontal regions that were notably absent in the AO condition.

A qualitative illustration of the pattern of flow rates for the top 10 mean values is shown in Fig. 3C (ranked based on the average value of |*τi*→*j*|). What is highlighted in these figures is that after observational practice, there is more involvement from the left frontal and parietal regions, as compared to M1-tDCS, and more transmission of information between the two hemispheres.

### 3.4 Statistical comparison of effective connectivity patterns

We performed a statistical comparison of the normalized information flow rates for each group, to determine if the information flow rate distributions were different. To do this, we calculated |τi→j| values across participants for each of the 90 unique pairs derived from 10 source locations. This causal connectivity analysis resulted in two distinct statistical distributions as shown in Fig. 4A, that based on a non-parametric Kruskal-Wallis (KW) test, showed a significant difference in the distribution of |τi→j| values across the two groups (*p <* 0.008). A further sub-analysis was conducted on the distribution, where we generated 100,000 sub-samples from a random sample of five individuals from each group and compared the coefficient of variation (COV), skewness, and kurtosis of |τi→j | distribution as shown in Fig. 4B. These plots highlight that active and passive stimulation are characterized by unique patterns of information flow rates.

**Fig. 4.**
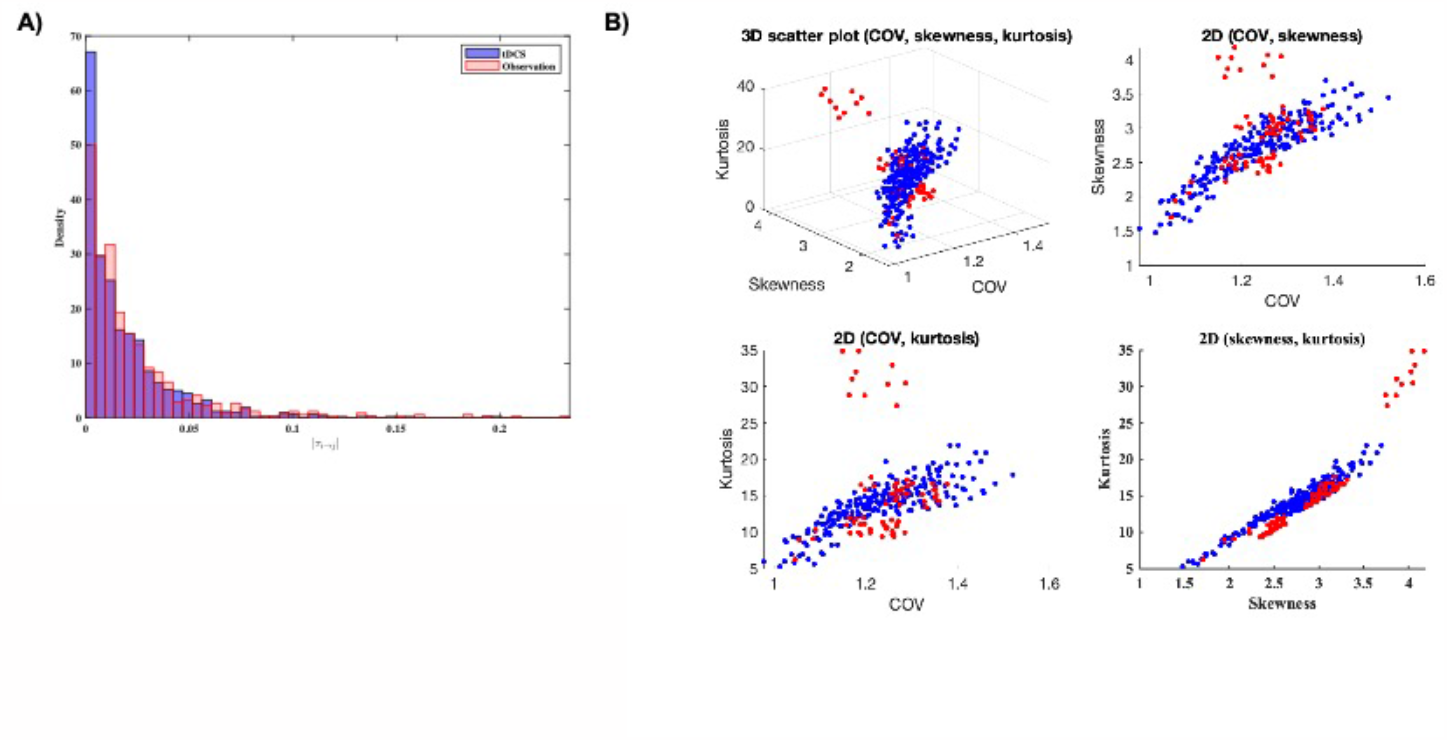
**(A)** Probability density histograms of the |τi→j | values for the M1-tDCS (blue) and observation (rose) groups. **(B)** Statistics of the |τi→j | distributions of each group. Top-left: A 3D scatter plot showing the relationship between COV, skewness, and kurtosis for tDCS group in blue and observation group in red. Top-right, bottom-left and bottom-right presenting 2D relationship between COV, skewness and kurtosis.

## 4 Discussion

In this study, we investigated effective connectivity of resting state EEG source data underlying skill acquisition of a complex, bimanual task as a function of two different types of stimulation – i.e. direct motor stimulation via M1-tDCS or indirect stimulation via AO. Overall, our data indicate that each intervention primes the brain through distinct network dynamics. M1-tDCS induced bi-directional changes in both the localized and more widespread brain regions whereas AO induced more targeted changes in motor and non-motor regions that are associated with the action-perception network. Notably, bimanual juggling accuracy was significantly higher following AO compared with M1-tDCS.

Contrary to our prediction related to AO, we found that the strongest connection associated with this condition was from LP to RP followed by bidirectional information transfer from RO and RP. It is well established that the parietal cortex has a central role in representing and interpreting actions (Chong et al, 2008; Hamilton & Grafton, 2006), with activity in the parietal regions occurring very early following the onset of movement observation (Virji-Babul et al., 2009). Furthermore, Urgen and Orban (2021) have recently proposed that the parietal cortex has a unique role in coding the immediate goal of the observable action. While these previous studies have shown the strongest areas of activation, we know little about the primary drivers of this activity. Our data show that information flow between left and right parietal regions and right parietal and occipital regions play a critical role not only in visual-perceptual processing, but also in contributing to a strong network involved in processing complex movements during action observation.

Among the top 5 connections in the AO group, LF (the left primary motor region) had strong information transfer to LT. Of particular note is that LT is a significant node within this network, with bi-directional information exchange occurring from frontal, motor, parietal and occipital regions. We have previously shown that during the observation of a right-handed, goal-directed reaching movement in a live model, the first brain area to be activated was the right temporal region (Virji-Babul et al., 2009) followed by activity in the sensorimotor and parietal regions, suggesting that this discrimination between self and other may be mediated by early interactions between the temporal regions and the sensorimotor regions. This may be an essential link between perception and action. It has been suggested that the temporal cortex interacts with premotor and parietal cortex particularly during imitation, by integrating visual input with reafferent copies of the imitated action. Recently, Pitcher and Ungerleider (2021) proposed a visual pathway from the visual cortex to the superior temporal sulcus (STS) to understand and interpret complex actions. While EEG does not have the spatial resolution for the fine-grained analysis of specific brain regions, our findings from information flow provide further evidence of the important role that the LT region plays in coding and integrating information from observation of a novel, complex task.

Our second prediction was based on past research findings showing that the effects of tDCS are not simply limited to regional effects but rather have widespread effects across the brain (Calzolari et al., 2023; Lang et al., 2005). Our results add to this literature by providing new information about the direction and strength of these distributed effects. Effective connectivity following direct stimulation of M1 was characterized by strong bilateral information exchange between the (R) occipital region and the (R) temporal regions. More aligned with our prediction, there was strong information exchange from both (L) and (R) frontal regions to (L) occipital and (L) temporal regions respectively as well as short-range, inter-hemispheric, bidirectional information transfer between the (L) and (R) motor regions. These results reveal that tDCS not only extends beyond the site of stimulation as reported previously, but that the strongest bidirectional information exchange occurs between right occipital and right temporal regions as well as long-range information flow from left frontal to left occipital regions. In addition, there is evidence of bidirectional, inter-hemispheric connections between left and right frontal regions that are absent in the AO condition. These results indicate that effective connectivity following M1-tDCS has a distinct pattern of activation that involves strong occipital-temporal information exchange and that left M1 stimulation induces motor activation across both hemispheres.

Our findings indicate that both AO and tDCS serve as tools for neural priming (e.g., Mark et al., 2023) and that they do so through different and distinct mechanisms. While both interventions demonstrated a clear impact on the plasticity of resting state effective connectivity, the AO group exclusively showed an enhancement in juggling accuracy. This result suggests that AO primes the brain in a more task-specific and goal-directed manner by stimulating networks both within the traditional mirror neuron network as well as broader visual-perceptual processes.

Conversely, tDCS likely primes the brain in a more generalized manner in preparation or readiness for a variety of different goals and behaviours. This preliminary study had a small sample size and further studies with larger samples are needed to validate the results.

## References

Alhajri, N., Hodges, N. J., Zwicker, J. G., & Virji-Babul, N. (2018). Mu Suppression Is Sensitive to Observational Practice but Results in Different Patterns of Activation in Comparison with Physical Practice. Neural Plasticity, 2018, e8309483. 10.1155/2018/8309483

Berchicci, M., Quinzi, F., Dainese, A., & Di Russo, F. (2017). Time-source of neural plasticity in complex bimanual coordinative tasks: Juggling. Behavioural Brain Research, 328, 87–94. 10.1016/j.bbr.2017.04.011

Calvo-Merino, B., Grèzes, J., Glaser, D. E., Passingham, R. E., & Haggard, P. (2006). Seeing or Doing? Influence of Visual and Motor Familiarity in Action Observation. Current Biology, 16(19), 1905–1910. 10.1016/j.cub.2006.07.065

Calzolari, S., Jalali, R., & Fernández-Espejo, D. (2023). Characterising stationary and dynamic effective connectivity changes in the motor network during and after tDCS. NeuroImage, 269, 119915. 10.1016/j.neuroimage.2023.119915

Ehsani, F., Bakhtiary, A. H., Jaberzadeh, S., Talimkhani, A., & Hajihasani, A. (2016). Differential effects of primary motor cortex and cerebellar transcranial direct current stimulation on motor learning in healthy individuals: A randomized double-blind sham-controlled study. Neuroscience Research, 112, 10–19. 10.1016/j.neures.2016.06.003

Friston, K. J. (2011). Functional and Effective Connectivity: A Review. Brain Connectivity, 1(1), 13–36. 10.1089/brain.2011.0008

Goodwill, A., Reynolds, J., Daly, R., & Kidgell, D. (2013). Formation of cortical plasticity in older adults following tDCS and motor training. Frontiers in Aging Neuroscience, 5. https://www.frontiersin.org/articles/10.3389/fnagi.2013.00087

Grèzes, J., Armony, J. L., Rowe, J., & Passingham, R. E. (2003). Activations related to “mirror” and “canonical” neurones in the human brain: An fMRI study. NeuroImage, 18(4), 928–937. 10.1016/S1053-8119(03)00042-9

Hamilton, A. F. de C., & Grafton, S. T. (2006). Goal representation in human anterior intraparietal sulcus. The Journal of Neuroscience: The Official Journal of the Society for Neuroscience, 26(4), 1133–1137. 10.1523/JNEUROSCI.4551-05.2006

Hodges, N. J. (2017). Observations on Action-Observation Research: An Autobiographical Retrospective Across the Past Two Decades. Kinesiology Review, 6(3), 240–260. 10.1123/kr.2017-0016

Hodges, N. J., & Coppola, T. (2015). What we think we learn from watching others: The moderating role of ability on perceptions of learning from observation. Psychological Research, 79(4), 609–620. 10.1007/s00426-014-0588-y

Hodges, N. J., Williams, A. M., Hayes, S. J., & Breslin, G. (2007). What is modelled during observational learning? Journal of Sports Sciences, 25(5), 531–545. 10.1080/02640410600946860

Hristopulos, D. T., Arif Babul, Babul, S., Brucar, L. R., & Virji-Babul, N. (2019). Disrupted Information Flow in Resting-State in Adolescents With Sports Related Concussion. Frontiers in Human Neuroscience, 13, 419. 10.3389/fnhum.2019.00419

Karok, S., & Witney, A. G. (2013). Enhanced Motor Learning Following Task-Concurrent Dual Transcranial Direct Current Stimulation. PLOS ONE, 8(12), e85693. 10.1371/journal.pone.0085693

Lang, N., Siebner, H. R., Ward, N. S., Lee, L., Nitsche, M. A., Paulus, W., Rothwell, J. C., Lemon, R. N., & Frackowiak, R. S. (2005). How does transcranial DC stimulation of the primary motor cortex alter regional neuronal activity in the human brain? The European Journal of Neuroscience, 22(2), 495–504. 10.1111/j.1460-9568.2005.04233.x

Liang, X. S. (2013). The Liang-Kleeman Information Flow: Theory and Applications. Entropy, 15(1), Article 1. 10.3390/e15010327

Liang, X. S. (2014). Unraveling the cause-effect relation between time series. Physical Review E, 90(5), 052150. 10.1103/PhysRevE.90.052150

Mark, J. I., Ryan, H., Fabian, K., DeMarco, K., Lewek, M. D., & Cassidy, J. M. (2023). Aerobic exercise and action observation priming modulate functional connectivity. PLOS ONE, 18(4), e0283975. 10.1371/journal.pone.0283975

Nitsche, M. A., & Paulus, W. (2000). Excitability changes induced in the human motor cortex by weak transcranial direct current stimulation. The Journal of Physiology, 527(Pt 3), 633–639. 10.1111/j.1469-7793.2000.t01-1-00633.x

Nitsche, M. A., & Paulus, W. (2001). Sustained excitability elevations induced by transcranial DC motor cortex stimulation in humans. Neurology, 57(10), 1899–1901. 10.1212/WNL.57.10.1899

Oldfield, R. C. (1971). The assessment and analysis of handedness: The Edinburgh inventory. Neuropsychologia, 9(1), 97–113. 10.1016/0028-3932(71)90067-4

Pitcher, D., & Ungerleider, L. G. (2021). Evidence for a Third Visual Pathway Specialized for Social Perception. Trends in Cognitive Sciences, 25(2), 100–110. 10.1016/j.tics.2020.11.006

Pixa, N. H., & Pollok, B. (2018). Effects of tDCS on Bimanual Motor Skills: A Brief Review. Frontiers in Behavioral Neuroscience, 12. https://www.frontiersin.org/articles/10.3389/fnbeh.2018.00063

Ramsey, R., Kaplan, D. M., & Cross, E. S. (2021). Watch and Learn: The Cognitive Neuroscience of Learning from Others’ Actions. Trends in Neurosciences, 44(6), 478–491. 10.1016/j.tins.2021.01.007

Rumpf, J.-J., Wegscheider, M., Hinselmann, K., Fricke, C., King, B. R., Weise, D., Klann, J., Binkofski, F., Buccino, G., Karni, A., Doyon, J., & Classen, J. (2017). Enhancement of motor consolidation by post-training transcranial direct current stimulation in older people. Neurobiology of Aging, 49, 1–8. 10.1016/j.neurobiolaging.2016.09.003

Stagg, C. J., Jayaram, G., Pastor, D., Kincses, Z. T., Matthews, P. M., & Johansen-Berg, H. (2011). Polarity and timing-dependent effects of transcranial direct current stimulation in explicit motor learning. Neuropsychologia, 49(5), 800–804. 10.1016/j.neuropsychologia.2011.02.009

Tadel, F., Baillet, S., Mosher, J. C., Pantazis, D., & Leahy, R. M. (2011). Brainstorm: A User-Friendly Application for MEG/EEG Analysis. Computational Intelligence and Neuroscience, 2011, e879716. 10.1155/2011/879716

Urgen, B. A., & Orban, G. A. (2021). The unique role of parietal cortex in action observation: Functional organization for communicative and manipulative actions. NeuroImage, 237, 118220. 10.1016/j.neuroimage.2021.118220

Virji-Babul, N., Moiseev, A., Cheung, T., Weeks, D., Cheyne, D., & Ribary, U. (2009). Spatial-temporal dynamics of cortical activity underlying reaching and grasping. Human Brain Mapping, 31(1), 160–171. 10.1002/hbm.20853

Zentgraf, K., & Munzert, J. (2009). Effects of attentional-focus instructions on movement kinematics. Psychology of Sport and Exercise, 10(5), 520–525. 10.1016/j.psychsport.2009.01.006

